# Endocervical regulatory T cells are associated with decreased genital inflammation and lower HIV target cell abundance

**DOI:** 10.1101/2021.03.28.437432

**Authors:** Aloysious Ssemaganda, Francois Cholette, Michelle Perner, Cheli Kambaran, Wendy Adhiambo, Peter M. Wambugu, Henok Gebrebrhan, Amy Lee, Faisal Nuhu, Ruth S. Mwatelah, Naima Jahan, Tosin E. Omole, Tabitha Wanjiru, Apollo Gitau, Joshua Kimani, Lyle R. McKinnon

**Affiliations:** Department of Medical Microbiology and Infectious Diseases, University of Manitoba, Winnipeg, Canada; JC Wilt Infectious Diseases Research Centre, Public Health Agency of Canada, Winnipeg, Canada; Department of Medical Microbiology, University of Nairobi, Nairobi, Kenya; Centre for the AIDS Programme of Research in South Africa (CAPRISA), Durban, South Africa

## Abstract

Regulatory T cells (Tregs) play important roles in tissue homeostasis, but few studies have investigated tissue Tregs in the context of genital inflammation, HIV target cell density, and vaginal microbiota in humans. In women from Nairobi (n=64), the proportion of CD4+ CD25+ CD127^low^ Tregs in the endocervix correlated with those in blood (r=0.31, p=0.01), with a higher Treg frequency observed in the endocervix (median 3.8 vs 2.0%, p<0.0001). Most Tregs expressed FoxP3 in both compartments, and CTLA-4 expression was higher on endocervical Tregs compared to blood (median 50.8 vs 6.0%, p<0.0001). More than half (34/62, 55%) of participants displayed a non-*Lactobacillus* dominant vaginal microbiota, which was not associated with endocervical Tregs or CD4+ T cell abundance. In a multivariable linear regression, endocervical Treg proportions were inversely associated with the number of elevated pro-inflammatory cytokines (p=0.03). Inverse Treg associations were also observed for specific cytokines including IL-1β, G-CSF, Eotaxin, IL-1RA, IL-8, and MIP-1 β. Higher endocervical Treg proportions were associated with lower abundance of endocervical CD4+ T cells (0.30 log_10_ CD4+ T cells per log_10_ Treg, p=0.00028), with a similar trend for Th17 cells (p=0.09). Selectively increasing endocervical Tregs may represent a pathway to reduce genital tract inflammation in women.

## Introduction

Genital inflammation defined by elevated levels of proinflammatory cytokines in mucosal secretions has been associated with ~3-fold increased HIV acquisition and diminished topical PrEP efficacy in women^1,2^. Additional work by our group has demonstrated that elevated cervicovaginal cytokines coincide with higher numbers of HIV target cells and a decrease in proteins associated with mucosal barrier function, providing a potential mechanism linking genital inflammation with HIV acquisition^3^. In this model, inflammatory cytokines recruit immune cells, which in turn secrete proteins that contribute toward mucosal tissue damage and further cytokine release, perpetuating a cycle of impaired barrier function and more target cell recruitment for HIV to establish a localized infection^4^. To disrupt this cycle, it is critical to understand the mechanisms that regulate inflammation and its pathogenic effects, which might contribute to interventions that can reduce the risk of HIV acquisition. While more attention has been directed toward the activation and memory status of CD4+ T cell subsets such as Th17 cells in the context of tissue inflammation in the cervix^5–9^, little is known about regulatory T cells (Tregs) in this tissue, and the roles they might play in the reducing inflammation and maintaining tissue homeostasis.

Tregs play a vital role in limiting excess immune responses and maintaining tissue immune tolerance and homeostasis^10^. Tregs are generally characterized on the basis of their origin, with natural Treg (nTreg) developing in the thymus and induced Treg (iTreg) derived from conventional CD4+ T cells in blood^11,12^. Tregs are defined by their memory and activation status, their phenotype, defined by expression of soluble and cell surface markers linked to their function, such as CTLA-4, PD-1, TGF-β, IL-10, and IL-35, their antigen specificity, and their relationship to other T helper subsets^13^. While iTregs often predominate in tissues and mature through interactions with cytokines and commensal flora, the general consensus is the provenance of tissue Tregs is dependent on the tissue examined^14,15^. Therefore, the balance and distribution of Tregs could have important implications for inhibition of specific inflammatory processes in tissues.

Fewer studies have focused on endocervical Tregs, with most focusing on their role in pregnancy/fertility outcomes^16–18^. This is in part because Treg frequency may be influenced by sex hormones; with FoxP3 expression being increased in the ovulatory phase of the menstrual cycle^19^, to increase odds of implantation. At the onset of fetal development, FoxP3+ Tregs migrate from peripheral blood to the maternal/fetal interface and mediate immune suppression^19^. Three nTreg and iTreg subsets; CD25^hi^ FoxP3+, PD1^hi^ IL-10+, and TIGIT+ FoxP3^dim^ were observed in the decidua and these contribute to tolerance during pregnancy ^20^.

Taken together, these data suggest there are many knowledge gaps regarding Tregs and the genital inflammatory milieu. We hypothesized that increased frequency of cervicovaginal Tregs would correspond to a reduction of genital inflammation. We tested this hypothesis amongst female sex workers in Nairobi, correlating endocervical Tregs to inflammatory surrogates of HIV susceptibility and adverse reproductive health outcomes. We found that cervicovaginal Tregs were associated with decreased genital inflammation and reduced frequencies of HIV target cell abundance, suggesting that increasing this cell population may represent a mechanism for better control of genital inflammation.

## Results

### Participant characteristics

HIV uninfected women were enrolled from the Sex Workers Outreach Program (SWOP) clinic in Nairobi (n=64, **Table 1**). The median age was 28 years (IQR: 25-30), and median time in sex work was 3 years (IQR: 2-6). Most (86%) reported sex work as their primary mode of income, and nearly three-quarters (72%) had completed secondary or tertiary education. Participants reported a median of one sex act with both repeat (IQR: 0-2) and casual clients (IQR: 0-3) in the week prior to enrolment. Condom use was quite common with casual and repeat clients, with >90% reporting always using condoms with these types of sexual partners. Just under half of participants (42%) reported having a husband or boyfriend, and ‘always’ condom use was much lower in this context (15%). Over half (53%) reported vaginal douching, and 67% reported use of enhancing substances during sex. Prior PrEP access was relatively common at 55% (35/64), and approximately one-quarter (17/64, 27%) had reported being previously diagnosed with a sexually transmitted infection. More than half (56%) reported have sex while under the influence of alcohol. Nearly half of participants (31/64, 48%) reported use of any form of contraception, with depot medroxyprogesterone acetate (DMPA) and implants being most commonly reported. The median number of pregnancies was 2 (IQR, 1-3) and 20% of the women reported a history of still birth, spontaneous abortion or miscarriage.

**Table 1.** Behavioral, reproductive and demographic characteristics of study participants

| **Variable** | **Frequency** |
| --- | --- |
| Age, median (IQR) | 28 (25-30) |
| Duration of sex work in years, median (IQR) | 3 (2-6) |
| Education, secondary school (or greater) completed | 46/64 (72%) |
| Earn most money from sex work | 55/64 (86%) |
| Have husband or boyfriend | 27/64 (42%) |
| Most recent sex, days ago, median (IQR) | 2 (1-7) |
| Sex acts with casual clients last week, median (IQR) | 1 (0-3) |
| Sex acts with repeat clients last week, median (IQR) | 1 (0-2) |
| Practice vaginal douching | 34/64 (53%) |
| Use enhancing substances during sex | 43/64 (67%) |
| Ever taken pre-exposure prophylaxis | 35/64 (55%) |
| Have sex during menses with casual clients | 20/64 (31%) |
| Always use condoms with casual clients | 51/52 (98%) |
| Always use condoms with repeat clients | 53/57 (93%) |
| Always use condoms with boyfriends/husbands | 4/27 (15%) |
| Number of pregnancies, median (IQR) | 2 (1-3) |
| Pregnancy resulting in still birth or miscarriage | 13/64 (20%) |
| Currently on contraceptives [DMPA (n=13), implant (n = 10), oral pills (n = 5)] | 31/64 (48%) |
| Menstrual cycle phase at sample visit, 16-30 days from start of menses (of those with regular cycles) | 22/57 (39%) |
| Ever diagnosed with a sexually transmitted infection | 17/64 (27%) |
| Ever exposed to physical violence | 16/64 (25%) |
| Ever exposed to sexual violence | 7/64 (11%) |
| Ever use recreational drugs (non-injection) | 17/64 (27%) |
| Ever have sex under influence of alcohol | 36/64 (56%) |

### Treg characterization in the female reproductive tract and blood

Tregs were defined by flow cytometry as live, single CD3+ CD4+ T cells expressing CD25 but not CD127 **(Fig. 1A).** Tregs were then gated separately to measure Treg-specific FoxP3+ and CTLA-4 expression. Th17 cells were characterized as CD4+ T cells expressing both CD161 and CCR6, as previously described **(Fig. 1B)**^21^. Most endocervical and peripheral Tregs expressed FoxP3 (median 45.3%, IQR: 27.1 to 60.0; and 45.3%, IQR: 25.9 to 58.8, respectively), with no difference between compartments (p=0.4, **Fig. 1C**). Endocervical Tregs expressed higher levels of CTLA-4 compared to blood (50.8%, IQR: 32.1 to 71.3 vs 6.04%, IQR: 2.2 to 10.7; p<0.0001, **Fig. 1D**). Treg frequency was higher in the cervix compared to blood (median 3.8%, IQR: 2.2 to 7.1 vs 2.0%, IQR 1.3 to 2.9; p<0.0001, **Fig. 1E**), but the proportion of Tregs in the endocervix correlated with those in blood (r=0.31, p=0.01, **Fig. 1F**).

**Figure 1.**
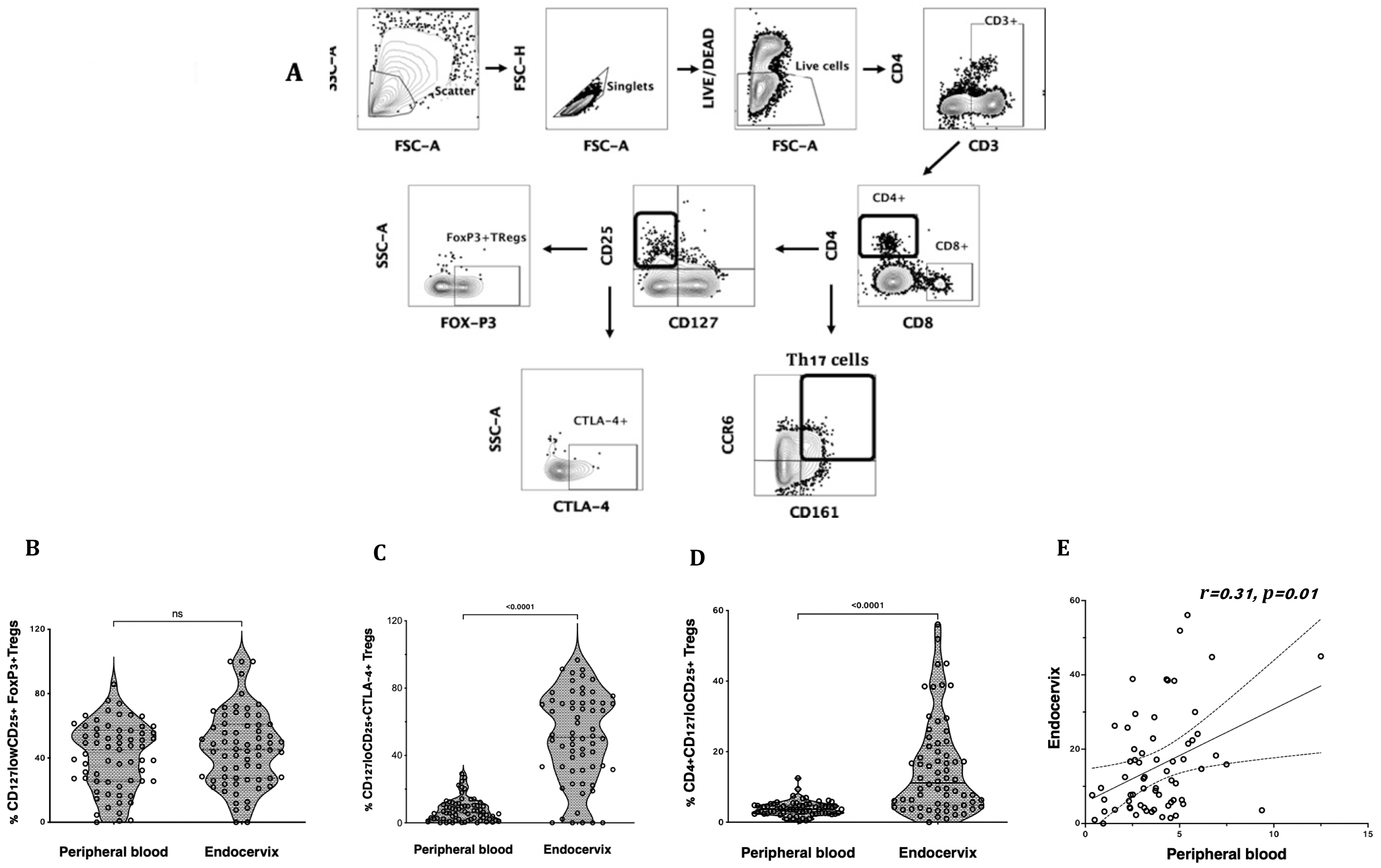
Treg and Th17 phenotyping in blood and endocervix. (A) Representative flow cytometry plots showing the gating strategy used to define Tregs as CD4+ T cells expressing CD25 and low CD127. These were then gated for FoxP3 and CTLA-4 expression. Th17 cells were defined by the expression of CD161 and CCR6. (B) Proportion of CD25+CD127lo FoxP3+ (C) CD25+CD127lo CTLA-4+ and (D) CD25+CD127lo Tregs in paired blood and cervicovaginal specimen (n=68) (D) CD25+CD127lo Tregs in the endocervix correlated with peripheral blood Tregs. ns: non-significant.

### Endocervical Treg associations with behavioral and reproductive variables

We next correlated Tregs with sexual behavioural, demographic and reproductive health variables. Endocervical Tregs correlated inversely with duration of sex work (r=-0.277, p=0.027, **Table 2**). The frequency of sex with repeat (r= −0.35, p=0.005) but not casual (r=0.038, p=0.766) sex partners correlated inversely with cervical Treg. History of stillbirth, miscarriage or spontaneous abortion was associated with lower frequencies of cervical Tregs (p=0.043). A trend toward lower Tregs was observed amongst women who used DMPA for contraception, compared to other methods, while the use of oral pills as contraception trended toward more endocervical Tregs (ANOVA p=0.107). No other sociodemographic, clinical or behavioural associations with Tregs were observed.

**Table 2.** Correlation of cervicovaginal Tregs with sexual behavioral and reproductive variables

| **Variable** | **Spearman rho** | **P value** |
| --- | --- | --- |
| Duration of sex work | -0.277 | 0.027 |
| No. of sex acts with repeat clients in the last week | -0.349 | 0.005 |
| No. of sex acts with casual clients in the last week | 0.038 | 0.766 |
| History of stillbirth, miscarriage or spontaneous abortion | -0.26 | 0.043 |
| Number of pregnancies | -0.145 | 0.254 |

### Cervicovaginal microbiome profiles and associations with Tregs

The relative abundance of cervicovaginal microbial communities was characterized by 16S rRNA sequencing of menstrual cup cell pellets. A majority of participants had a non-*Lactobacillus* dominant profile (n = 34, 55%, **Fig. 2A**). Unsupervised hierarchical clustering differentiated the OTUs into three clusters, with *Lactobacillus* dominant as branch 1 (n = 28, 45%), a mixture *Sneathia, Prevotella*, and *Gardnerella* as branch 2 (n = 25, 40%), and a predominance of *Gardnerella* as branch 3 (n = 9, 15%). The most common *Lactobacillus* species in branch 1 was *L. iners*. The alpha diversity of branches 2 and 3 were significantly higher than branch 1 (not shown). No associations were observed between vaginal microbial cluster or diversity with any endocervical cell variable, including Treg frequency and absolute CD4+ T cell abundance **(Fig. 2B & C)**.

**Figure 2.**
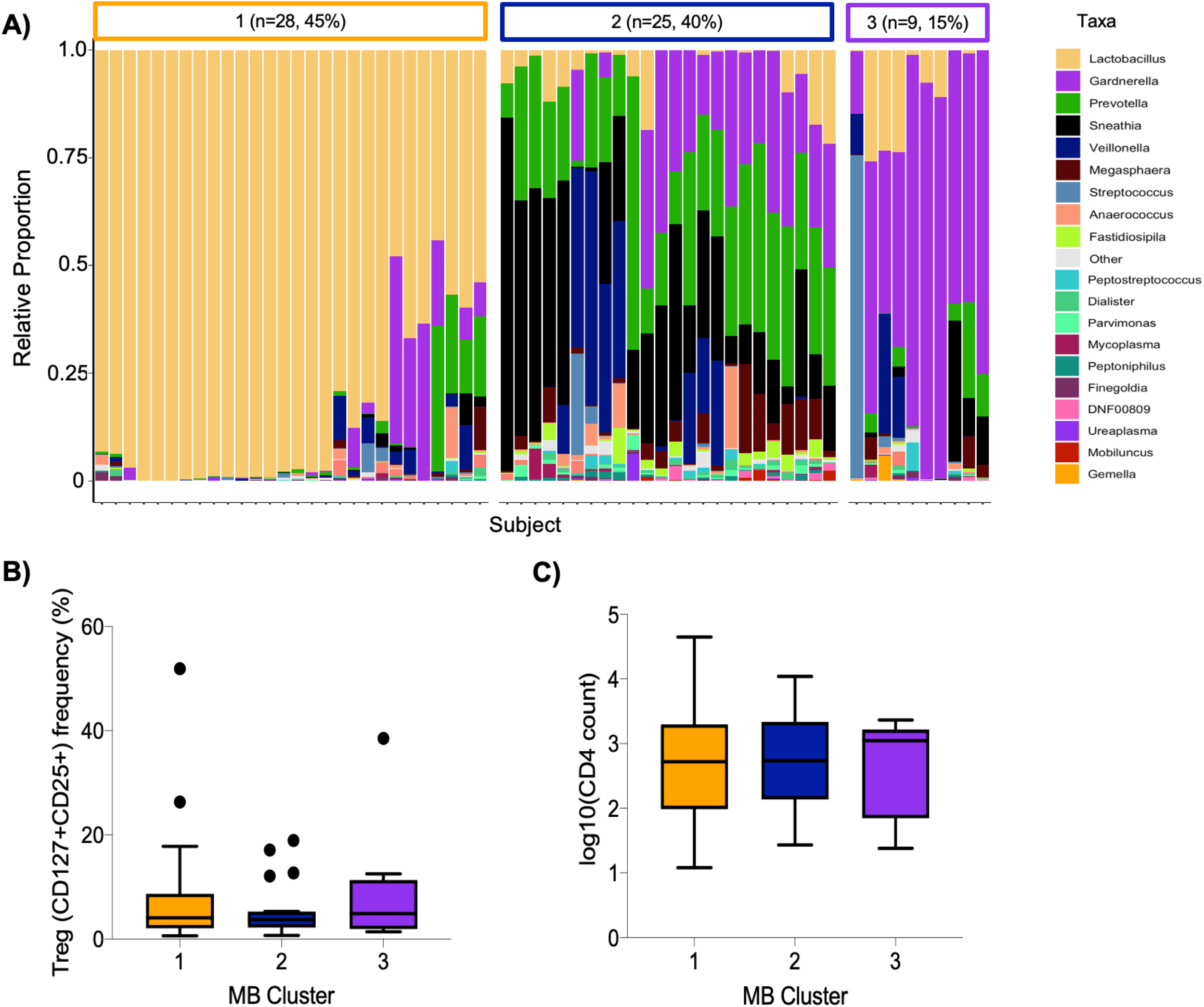
Cervicovaginal microbial profiles amongst female sex workers from Nairobi, Kenya (n = 62). (A) Unsupervised clustering of relative abundance of microbial communities summarized by taxa with 45% being *Lactobacillus* dominant, 40% a mixture *Sneathia, Prevotella*, and *Gardnerella* and 15% predominantly *Gardnerella spp*. B) Association of microbial clusters 1, 2 and 3 with Treg frequency and (C) absolute CD4 T cell count.

### Endocervical Tregs and genital inflammation

We have previously defined genital inflammation based on the number of pro-inflammatory cytokines in the upper quartile^1,2^. In this study, we quantified cytokine concentrations in cervical secretions collected via menstrual cups and found that approximately one-quarter (24%, n=15/62) had ≥ 3 pro-inflammatory cytokines in the upper quartile. We modeled number of elevated cytokines as a continuous variable. Although the number of elevated genital cytokines was not associated with endocervical Tregs prior to adjustment, an inverse correlation was observed in an adjusted linear regression model with endocervical Treg as the outcome (beta = −0.07, 95% CI: −0.13 to −0.01, p = 0.028, **Table 3**). To better understand which cytokines in the multiplex panel were most associated with endocervical Treg, we correlated each cytokine as a log_10_ pg/ml with normalized Treg frequency; six cytokines were inversely associated with Treg in a multivariable analysis (**Table 3**), the strongest of which were the pro-inflammatory cytokine IL-1β (p = 0.00019) and its related antagonist IL-1RA (p = 0.005), the growth factor G-CSF (p = 0.00022), and the chemokines IL-8 (p = 0.02), MIP-1β (p = 0.031) and Eotaxin (p = 0.002). Clear step-wise decreases in concentration of each of these cytokines was observed by endocervical Treg tertile (**Figure 3).**

**Table 3.**
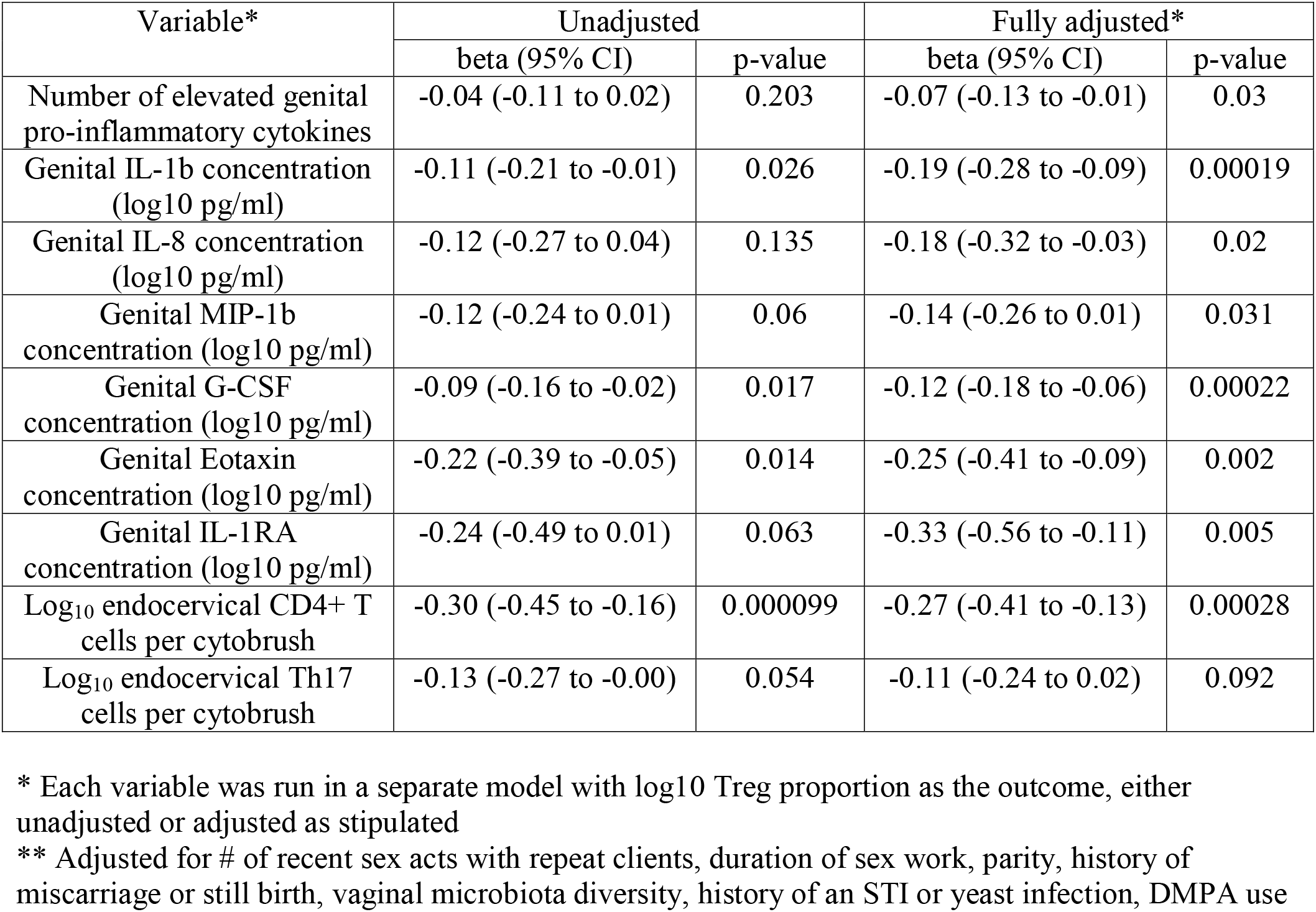
Multivariable models of endocervical Treg frequency.

| Variable* | Unadjusted |  | Fully adjusted* |  |
| --- | --- | --- | --- | --- |
|  | beta (95% CI) | p-value | beta (95% CI) | p-value |
| Number of elevated genital pro-inflammatory cytokines | -0.04 (-0.11 to 0.02) | 0.203 | -0.07 (-0.13 to -0.01) | 0.03 |
| Genital IL-1b concentration (log10 pg/ml) | -0.11 (-0.21 to -0.01) | 0.026 | -0.19 (-0.28 to -0.09) | 0.00019 |
| Genital IL-8 concentration (log10 pg/ml) | -0.12 (-0.27 to 0.04) | 0.135 | -0.18 (-0.32 to -0.03) | 0.02 |
| Genital MIP-1b concentration (log10 pg/ml) | -0.12 (-0.24 to 0.01) | 0.06 | -0.14 (-0.26 to 0.01) | 0.031 |
| Genital G-CSF concentration (log10 pg/ml) | -0.09 (-0.16 to -0.02) | 0.017 | -0.12 (-0.18 to -0.06) | 0.00022 |
| Genital Eotaxin concentration (log10 pg/ml) | -0.22 (-0.39 to -0.05) | 0.014 | -0.25 (-0.41 to -0.09) | 0.002 |
| Genital IL-1RA concentration (log10 pg/ml) | -0.24 (-0.49 to 0.01) | 0.063 | -0.33 (-0.56 to -0.11) | 0.005 |
| Log<sub>10</sub> endocervical CD4+ T cells per cytobrush | -0.30 (-0.45 to -0.16) | 0.000099 | -0.27 (-0.41 to -0.13) | 0.00028 |
| Log<sub>10</sub> endocervical Th17 cells per cytobrush | -0.13 (-0.27 to -0.00) | 0.054 | -0.11 (-0.24 to 0.02) | 0.092 |

\* Each variable was run in a separate model with log10 Treg proportion as the outcome, either unadjusted or adjusted as stipulated
\*\* Adjusted for # of recent sex acts with repeat clients, duration of sex work, parity, history of miscarriage or still birth, vaginal microbiota diversity, history of an STI or yeast infection, DMPA use

**Figure 3.**
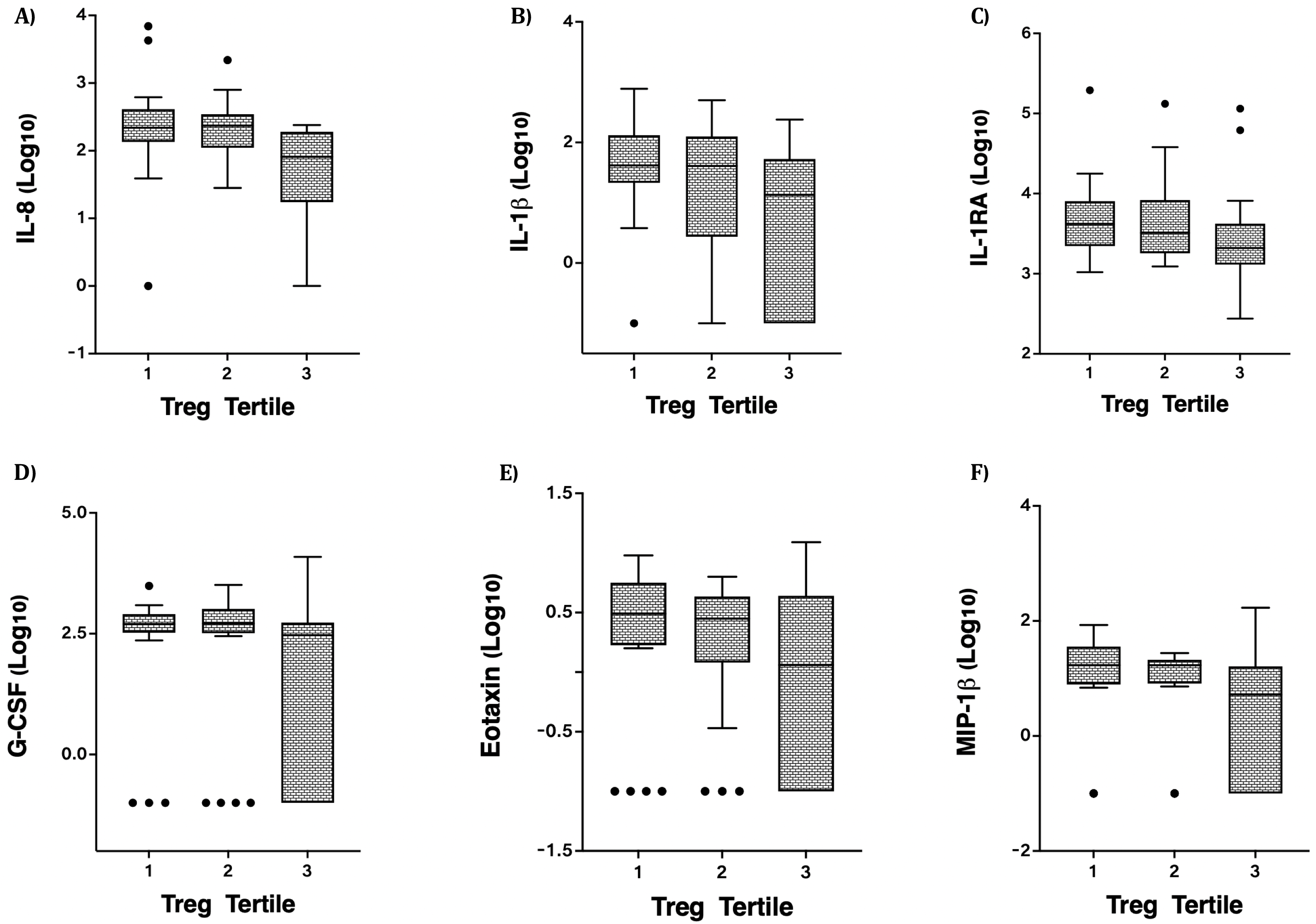
Correlation of the cervicovaginal Treg frequency with genital cytokine/chemokine concentrations. Concentrations of (A) IL-8 (B) IL-1β (C) IL-1RA (D) G-CSF, (E) Eotaxin and MIP-1β by endocervical Treg tertile. For statistical analysis, see Table 3.

### Endocervical Tregs and HIV target cells

We also measured associations between Tregs and other T cell subsets in the same endocervical samples, particularly cells that have been described as highly susceptible to HIV infection^22^. In the endocervix, higher Treg proportions were associated with lower abundance of CD3+ T cells (data not shown), CD4+ T cells, and Th17 cells defined by CCR6 and CD161 expression (**Figure 4)**. Endocervical CD4+ T cell abundance was inversely associated with endocervical Treg frequency in both unadjusted (p=0.000099) and adjusted models (beta = −0.27, 95% CI: −0.41 to −0.13, p=0.00028, **Table 3**). The mean number of endocervical CD4+ T cells declined step-wise, from 3.1 to 2.7 to 2.3 log_10_ CD4+ T cell abundance, a nearly 10-fold reduction in the number of CD4+ T cells from highest to lowest Treg tertile. Endocervical Th17 abundance was inversely correlated with endocervical Tregs in unadjusted models (p=0.054) and remained as a similar trend in multivariable analysis (p=0.092).

**Figure 4.**
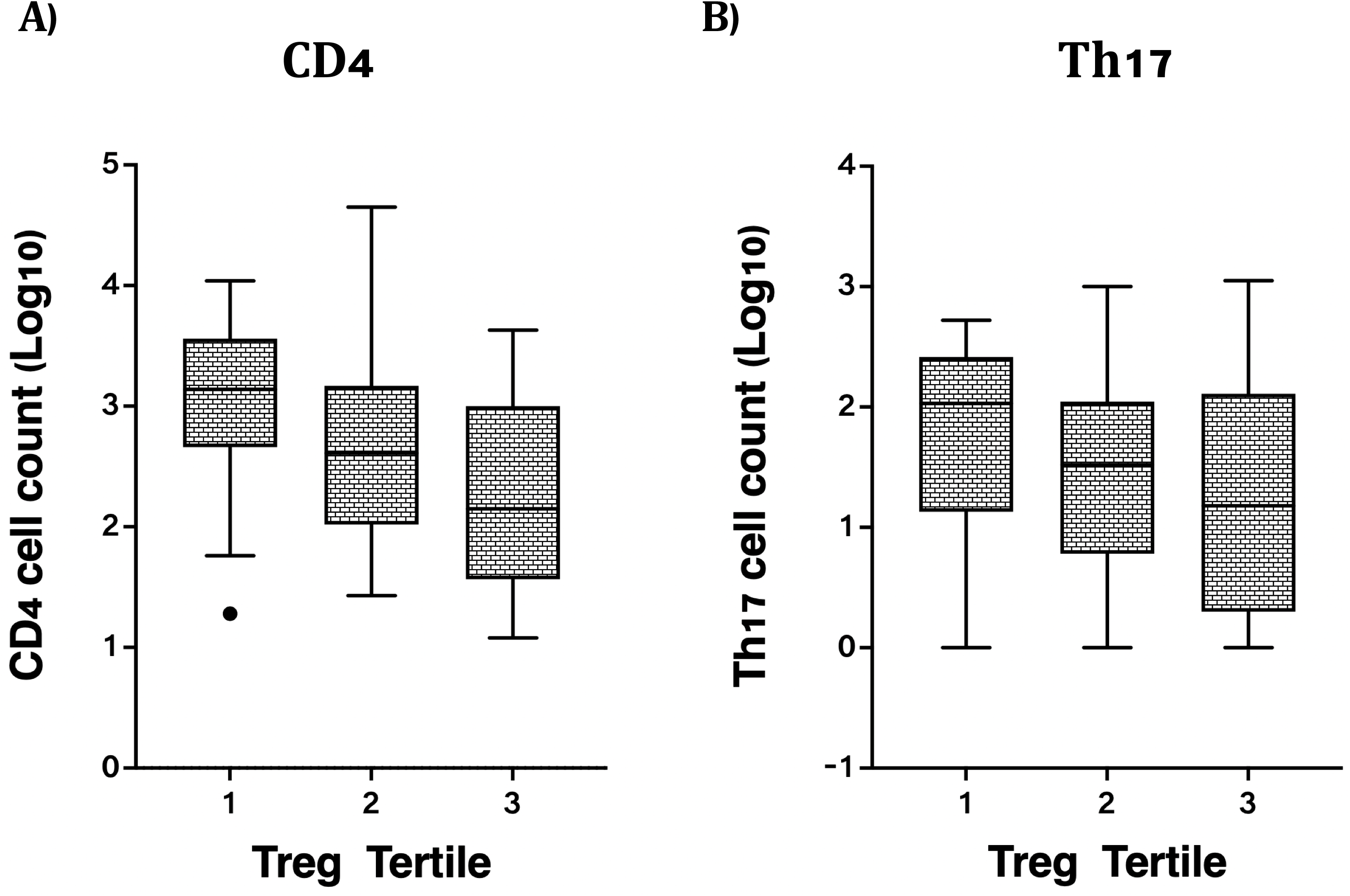
Correlation of cervicovaginal Treg frequencies with absolute counts of HIV target cells per cytobrush, including: (A) CD4 and (B) Th17 counts in the female genital tract. For statistical analysis, see Table 3.

## Discussion

Despite efforts to reduce HIV incidence using the currently available prevention, UNAIDS estimates that over 1.5 million new infections continue to occur annually, further underscoring the need to develop new tools meet UN targets to reduce HIV incidence by 90% by 2030^23^. Many of the biological factors implicated in HIV transmission have genital inflammation as a central feature^4^. While the causes and consequences of inflammation have been the focus of many studies, less attention has been paid to natural regulatory mechanisms that might restrain inflammatory responses despite exposure to inflammatory stimuli. Here we show that Tregs are more frequent in the endocervix than the blood, and display high expression of both FoxP3 and CTLA-4. Furthermore, their frequency correlates inversely with both the concentrations of inflammatory cytokines and the abundance of HIV target cells in the genital tract. While it is not possible to determine causal relationships in this cross-sectional study, one interpretation of these data is that increased Tregs in tissue are able to control the inflammatory responses generated by epithelial and/or immune cells in response to inflammatory stimuli. If confirmed, inducing Treg using interventions such as low dose IL-2 could reduce genital inflammation^24,25^, with important implications for preventing STIs and improving reproductive health.

Epidemiological associations of endocervical Tregs involved reproductive health variables, in particular a history of miscarriage or stillbirth. Although the study was not designed for this purpose, the inflammation-reducing potential of Treg could be investigated further, as inflammation has been associated with adverse reproductive health outcomes in addition to HIV^26^. Sexual intercourse with repeat but not casual clients was also associated inversely with endocervical Tregs. The reason for this difference is unclear and requires further investigation. We were unable to evaluate condom use, as this is reported to be quite high (>90% ‘always’ with both client types). Understanding natural variation in endocervical Tregs between women is an important further research objective.

Our findings add to the literature around reproductive tract Tregs outside of the context of reproductive variables. Similar to previous work, we find Treg are proportionally increased in the endocervix compared to the blood, by approximately two-fold^27^. Interestingly, with respect to pathogens, a balance is required between Tregs that protect against excess inflammation, facilitating protective immunity (as is the case with HSV-2, ^28,29^) versus those that can be co-opted to limit host immunity as a persistence strategy, as has been suggested for gonorrhea^30^. In the HIV acquisition context, the finding that women whose CD4+ T cell populations contained proportionally higher Treg had lower concentrations of inflammatory cytokines and HIV target cells suggests that Tregs may play a protective role by limiting HIV’s ability to cross the mucosal barrier and establish local infection, both thought to be critical events in transmission success. Experiments to test this hypothesis formally are needed.

While Tregs may protect against HIV by limiting inflammation, they also regulate frequencies of Th17 cells^31^, which we and others have shown to be preferential targets for HIV in the mucosa^5,7,8^. In the present study, we observed an inverse trend between endocervical Treg and Th17; a larger, prospective study that induces Treg would be needed to determine the impact of increasing Treg on Th17 cells in this tissue.

We did not observe any associations between endocervical Treg and vaginal microbiota. While this needs to be confirmed in a larger, prospective study, one explanation is that Treg regulate the inflammatory response to non-optimal, bacterial vaginosis (BV) associated bacteria like *Sneathia*, *Gardnerella,* and *Prevotella*, rather than their presence in the mucosa. This could be an important consideration, as bacterial vaginosis, a condition marked by depletion of vaginal health associated *Lactobacillus*, is associated with increased risk of STI acquisition^32–35^, reproductive complications^36–38^, and is highly recalcitrant to existing treatments^39^, so modulating the detrimental effects of bacteria could be an alternative or additional approach to minimize the impact of BV. Similar to what we have previously shown in a cohort of adolescent girls and young women in Mombasa, Kenya^40^, our current data also indicates that involvement in sex work might be associated with non-optimal vaginal microbiomes. Most women in SWOP displayed a non-*Lactobacillus* microbial profile characterized predominantly by *Sneathia, Prevotella*, and *Gardnerella*. Previous studies have shown that *Lactobacillus crispatus* induces an anti-inflammatory phenotype that included increased Tregs in the female reproductive tract (FRT)^41^. However, this could not be confirmed in this study due to the infrequency of these ‘optimal’ bacteria in our study population. These data underscore the need to improve vaginal and reproductive health in key populations at risk of HIV acquisition.

Our study had some important limitations. The cross-sectional design and modest sample size were sufficient to test associations between Treg and other variables including those related to inflammation, but cannot infer causation and provide limited opportunity to stratify the data extensively. Literature on tissue Tregs is expanding, and our flow cytometry panel was unable to fully phenotype emerging subsets, such as those co-expressing RORC, which regulate Th17 cells in the gut^42^, or that co-express GATA3, which are important in repairing fibrosis in the skin^43^. We did not collect ectocervical biopsies in this study, and Tregs in this tissue may differ from those recovered using endocervical cytobrushes.

In summary, our data provide important insights into mechanisms that may be critical in regulating inflammation and its pathogenic effects. Since the causes of genital inflammation are quite broad, with sources including sexual intercourse^40^, intravaginal practices, bacterial/viral sexually transmitted infections (STIs)^44,45^, and non-*Lactobacillus* vaginal microbiota associated with bacterial vaginosis^46^, one advantage of inducing regulatory mechanisms to reduce inflammation is that this regulation could protect against inflammation regardless of the cause. The strongest immune associations with endocervical Treg included overall CD4+ T cell abundance, which are believed to be required for HIV to establish itself in the mucosa, and a number of cytokines including those that attract immune cells to the site. Further exploration of associations between Treg and other immune cell subsets may be critical to understand precise mechanisms by which these effects are manifested.

These data suggest that inducing Tregs in the FRT might limit inflammation, and by extension, reduce HIV acquisition risk and improve reproductive health.

## Methods

### Study population and ethics statement

Female HIV sero-negative women (n=68) accessing care at Sex Workers Outreach Program (SWOP) clinic, Nairobi City, were enrolled in the study. The study was approved by the Kenyatta National Hospital - University of Nairobi Ethics and Research Committee. Prior to enrolment, written informed consent was obtained from all volunteers. A comprehensive questionnaire was then administered to establish each participant’s socio-demographic, reproductive and behavioural characteristics.

### Sample collection and processing

Cervicovaginal secretions were collected by insertion of a menstrual cup (SoftCup™) into the vagina during administration of the study questionnaire for at least 1 hour. On removal, the menstrual cup was transferred to a 50ml tube and kept on ice. In the laboratory, the Soft Cup secretions were resuspended in a 1:4 ratio in cold phosphate buffered saline (PBS), centrifuged, supernatant aliquoted in cryovials and the pellet resuspended in RNA-Later. Both the supernatant and pellet were stored − 80°C for cytokine and microbiome profiling respectively. Paired blood (10ml), in Acid Citrate Dextrose (ACD) tubes and cervical cytobrushes were collected from all enrolled volunteers for peripheral blood mononuclear cells (PBMC) and Cervical Mononuclear Cells (CMC) isolation respectively. Two cytobrushes were serially inserted into the endocervix, rotated through 360° transferred into a conical tube containing 3ml of PBS and placed on ice for processing within 2 hours of collection. In the laboratory, the tube containing cytobrushes was vigorously vortexed to dissociate the cells and mucus from the brushes, rinsed with R-10 medium (10% Foetal Bovine Serum (FBS) in RPMI containing penicillin and streptomycin) and filtered through 100-micron nylon cell strainer (Becton Dickinson) fitted into a 50 ml tube. Cells were washed twice by centrifugation at 1600rpm and pellet resuspended in washing buffer (PBS 2% FBS). PBMCs were isolated by Ficoll density separation and reconstituted in washing buffer. *Ex vivo* phenotyping of regulatory T cells (Tregs) and Th17 cells by flow cytometry was then performed on fresh CMC and PBMC.

### Flow cytometry

Cells were stained with viability exclusion dye, Fixable Aqua Live/Dead Stain (Life Technologies) in PBS at 4°C for 20 min. Cells were washed and surface stained for 30 min at 4°C with an optimized panel of fluorochrome labeled antibodies specific for the following antigens: CD3 Alexa-flour 700 (UCHT1), CD4 PerCP (OKT4), CD8 APC-Cy7 (RPA-T8), CCR6 BV421 (11A9), CD25 BB515 (M-A251), CD127 PE (A019D5), CD161 PE-eFluor 610 (HP-3G10) and CTLA-4 APC (L3D10). Cells were subsequently washed, permeabilized using the eBioscience fixation/ permeabilization solution (eBiosciences; San Diego, CA, USA) and stained with FoxP3 PE-Cy7 (PCH101) monoclonal antibody. The cells were washed again and resuspended in perm/wash buffer (eBioscience; San Diego, CA, USA). Data were acquired using an LSRII flow cytometer (BD Biosciences) and analyzed using FlowJo™ version 10.6.1 (Becton Dickinson Life Sciences).

### Cytokine/chemokine analysis

As previously described^2^, cytokine/chemokine concentrations were examined from 50μL of undiluted cervical secretions using the Bio-Plex Pro™ Human multiplex kit (IL-1β, IL-6, TNF-α, IL-8, IP-10, MCP-1, MIP-1α, MIP-1β, IL-1RA, G-CSF and Eotaxin; Bio-Rad Laboratories Inc.) following the manufacturer’s instructions. The plate was read using the Bio-Plex 200 Array instrument (Bio-Rad Laboratories) and data analysed using Bio-Plex Manager software version 6.1.1, to determine cytokine concentrations in pg/ml from the standard curve.

### Microbiome analysis

We performed 16S rRNA sequencing of bacterial DNA isolated from menstrual cup cell pellets according to Illumina’s 16S metagenomics sequencing library preparation guide using the MiSeq Reagent Kit v3 (600-cycle) and MiSeq sequencing platform according to the manufacturer’s instructions (https://support.illumina.com/documents/documentation/chemistry_documentation/16s/16s-metagenomic-library-prep-guide-15044223-b.pdf). DNA was extracted using the DNeasy Blood & Tissue kit (Qiagen, Hilden, Germany). Prior to DNA isolation, cell pellets were homogenized in approximately 200 μL of PBS using pre-filled silica bead tubes (VWR, Radnor, Pennsylvania) and a Bead Ruptor Elite homogenizer (OMNI International, Kennesaw, Georgia) using the following settings: 3 cycles of bead beating at 6 m/s for 45 seconds followed by a 2 min dwell period. PCR amplification verification and library validation was done using a TapeStation electrophoresis system and D1000 ScreenTape reagents (Agilent, Santa Clara, California). Libraries were quantified using a Qubit 4 Fluorometer and the Qubit dsDNA HS assay kit (ThermoFisher Scientific, Waltham, Massachusetts). The QIIME2 platform was used to process sequence data, perform quality control, and generate a table representing the number of reads that mapped to specific operational taxonomic units (OTUs).

### Statistical analysis

Descriptive statistics including medians and interquartile ranges, and proportions, were used to summarize continuous and categorical variables. These clinical, sociodemographic, behavioural and reproductive health variables were also correlated to the proportion of endocervical Treg and Th17, and the abundance of Treg, Th17, and CD4+ T cells, using Spearman rank correlation and Chi-squared tests. Vaginal microbiome data, using genera relative proportions, were grouped based on unsupervised hierarchical Euclidean clustering, and results were visualized with R (package ‘ggplot2’, Wickham, v3.3.2, 2016), and correlated to Treg proportions. Centered log ratio (CLR) transformations of proportional cellular data (Cell proportions and bacterial relative proportions) were performed in R (package ‘clr’, Pierrot, v0.1.2, 2019) before Spearman rank correlation calculations with PRISM (v9.0.0). Unmapped bacterial sequences were removed from analysis, and the lowest abundance 52 taxa were grouped into “other” for visualization (average=0.4±0.7% per sample).

Associations with other immune variables were tested using multivariable linear regression, with log_10_ transformed endocervical Treg proportions as the outcome. Separate models were used for each immune variable (number of cytokines, log_10_ concentration of genital cytokines, and log_10_ endocervical cell counts), adjusting for covariates associated with inflammation and/or Tregs, either in this study or in the literature. Models were adjusted for duration of sex work and not age, as the former was associated with endocervical Treg. Immune variables were graphed by Treg tertile for visualization purposes, but analysed as continuous variables. Graphing and visualization was performed using Prism version 9 (GraphPad Software, LLC), with additional analyses performed using R and SPSS v. 27.0.

## Acknowledgments

We thank all the study participants and the clinical staff at the SWOP City clinic in Nairobi, Kenya who made the study happen. We acknowledge the Canadian Institutes of Health Research CIHR PJT-148796) and the Manitoba Medical Service Foundation (MMSF 8-2019-06) for their generous funding support. LRM is supported by a CIHR New Investigator Award.

## Author Contributions

Study design and conceptualization: A.S., J.K., and L.R.M.

Performed the experiments: A.S., F.C., A.L., W.A., P.M.W., H.G., F.B.N., T.E.O., R.S.M., and N.J.

Analyzed the data: A.S., C.K., H.G., M.P., and L.R.M.

Wrote original manuscript: A.S. and L.R.M., with all authors contributed to manuscript editing and review.

Clinical specimen collection and administration of socio-behavioral and demographic questionnaire: T.W. and A.G.

Supervised clinical and/or experimental aspects of the study: J.K. and L.R.M.

## Disclosure

No conflicts of interest to declare.

## Notes

### Competing Interest Statement

The authors have declared no competing interest.

## References

1. Masson, L. et al. Genital inflammation and the risk of HIV acquisition in women. Clin. Infect. Dis. 61, 260–269 (2015).

2. McKinnon, L. R. et al. Genital inflammation undermines the effectiveness of tenofovir gel in preventing HIV acquisition in women. Nat. Med. 24, 491–496 (2018).

3. Arnold, K. B. et al. Increased levels of inflammatory cytokines in the female reproductive tract are associated with altered expression of proteases, mucosal barrier proteins, and an influx of HIV-susceptible target cells. Mucosal Immunol 9, 194–205 (2016).

4. Kaul, R. et al. Inflammation and HIV Transmission in Sub-Saharan Africa. Curr HIV/AIDS Rep 12, 216–222 (2015).

5. McKinnon, L. R. et al. Characterization of a human cervical CD4+ T cell subset coexpressing multiple markers of HIV susceptibility. J. Immunol. 187, 6032–6042 (2011).

6. McKinnon, L. R. et al. Early HIV-1 infection is associated with reduced frequencies of cervical Th17 cells. J. Acquir. Immune Defic. Syndr. 68, 6–12 (2015).

7. Stieh, D. J. et al. Th17 Cells Are Preferentially Infected Very Early after Vaginal Transmission of SIV in Macaques. Cell Host Microbe 19, 529–540 (2016).

8. Planas, D. et al. HIV-1 selectively targets gut-homing CCR6+CD4+ T cells via mTOR-dependent mechanisms. JCI Insight 2, 519 (2017).

9. Rodriguez-Garcia, M., Barr, F. D., Crist, S. G., Fahey, J. V. & Wira, C. R. Phenotype and susceptibility to HIV infection of CD4+ Th17 cells in the human female reproductive tract. Mucosal Immunol 7, 1375–1385 (2014).

10. Sakaguchi, S., Sakaguchi, N., Asano, M., Itoh, M. & Toda, M. Immunologic self-tolerance maintained by activated T cells expressing IL-2 receptor alpha-chains (CD25). Breakdown of a single mechanism of self-tolerance causes various autoimmune diseases. J. Immunol. 155, 1151–1164 (1995).

11. Caramalho, Í., Nunes-Cabaço, H., Foxall, R. B. & Sousa, A. E. Regulatory T-Cell Development in the Human Thymus. Front Immunol 6, 395 (2015).

12. Schmitt, E. G. & Williams, C. B. Generation and function of induced regulatory T cells. Front Immunol 4, 152 (2013).

13. Sakaguchi, S. et al. Regulatory T Cells and Human Disease. Annu Rev Immunol 38, 541–566 (2020).

14. Sharma, A. & Rudra, D. Emerging Functions of Regulatory T Cells in Tissue Homeostasis. Front Immunol 9, 883 (2018).

15. Panduro, M., Benoist, C. & Mathis, D. Tissue Tregs. Annu Rev Immunol 34, 609–633 (2016).

16. Robertson, S. A. & Moldenhauer, L. M. Immunological determinants of implantation success. Int. J. Dev. Biol. 58, 205–217 (2014).

17. de Barros, I. B. L. et al. “What do we know about regulatory T cells and endometriosis? A systematic review”. J. Reprod. Immunol. 120, 48–55 (2017).

18. Clark, D. A. & Chaouat, G. Regulatory T cells and reproduction: how do they do it? J. Reprod. Immunol. 96, 1–7 (2012).

19. Walecki, M. et al. Androgen receptor modulates Foxp3 expression in CD4+CD25+Foxp3+ regulatory T-cells. Mol. Biol. Cell 26, 2845–2857 (2015).

20. Salvany-Celades, M. et al. Three Types of Functional Regulatory T Cells Control T Cell Responses at the Human Maternal-Fetal Interface. Cell Rep 27, 2537–2547.e5 (2019).

21. Boily-Larouche, G. et al. CD161 identifies polyfunctional Th1/Th17 cells in the genital mucosa that are depleted in HIV-infected female sex workers from Nairobi, Kenya. Sci Rep 7, 11123–10 (2017).

22. McKinnon, L. R. & Kaul, R. Quality and quantity: mucosal CD4+ T cells and HIV susceptibility. Curr Opin HIV AIDS 7, 195–202 (2012).

23. Political Declaration on HIV and AIDS: On the Fast Track to Accelerating the Fight against HIV and to Ending the AIDS Epidemic by 2030: https://www.unaids.org/en/resources/documents/2016/2016-political-declaration-HIV-AIDS. at < https://www.unaids.org/en/resources/documents/2016/2016-political-declaration-HIV-AIDS>

24. Abbas, A. K., Trotta, E., R Simeonov, D., Marson, A. & Bluestone, J. A. Revisiting IL-2: Biology and therapeutic prospects. Sci Immunol 3, eaat1482 (2018).

25. Eskandari, S. K. et al. Regulatory T cells engineered with TCR signaling-responsive IL-2 nanogels suppress alloimmunity in sites of antigen encounter. Sci Transl Med 12, (2020).

26. Fettweis, J. M. et al. The vaginal microbiome and preterm birth. Nat. Med. 25, 1012–1021 (2019).

27. Iyer, S. S. et al. Characteristics of HIV target CD4 T cells collected using different sampling methods from the genital tract of HIV seronegative women. PLoS ONE 12, e0178193 (2017).

28. Soerens, A. G., Da Costa, A. & Lund, J. M. Regulatory T cells are essential to promote proper CD4 T-cell priming upon mucosal infection. Mucosal Immunol 9, 1395–1406 (2016).

29. Milman, N. et al. In Situ Detection of Regulatory T Cells in Human Genital Herpes Simplex Virus Type 2 (HSV-2) Reactivation and Their Influence on Spontaneous HSV-2 Reactivation. J. Infect. Dis. 214, 23–31 (2016).

30. Imarai, M. et al. Regulatory T cells are locally induced during intravaginal infection of mice with Neisseria gonorrhoeae. Infect. Immun. 76, 5456–5465 (2008).

31. Weaver, C. T. & Hatton, R. D. Interplay between the TH17 and TReg cell lineages: a (co-)evolutionary perspective. Nat. Rev. Immunol. 9, 883–889 (2009).

32. Atashili, J., Poole, C., Ndumbe, P. M., Adimora, A. A. & Smith, J. S. Bacterial vaginosis and HIV acquisition: a meta-analysis of published studies. AIDS 22, 1493–1501 (2008).

33. Gallo, M. F. et al. Bacterial vaginosis, gonorrhea, and chlamydial infection among women attending a sexually transmitted disease clinic: a longitudinal analysis of possible causal links. Ann Epidemiol 22, 213–220 (2012).

34. Gillet, E. et al. Bacterial vaginosis is associated with uterine cervical human papillomavirus infection: a meta-analysis. BMC Infect Dis 11, 10–9 (2011).

35. Cherpes, T. L., Meyn, L. A., Krohn, M. A., Lurie, J. G. & Hillier, S. L. Association between acquisition of herpes simplex virus type 2 in women and bacterial vaginosis. Clin. Infect. Dis. 37, 319–325 (2003).

36. Leitich, H. & Kiss, H. Asymptomatic bacterial vaginosis and intermediate flora as risk factors for adverse pregnancy outcome. Best Pract Res Clin Obstet Gynaecol 21, 375–390 (2007).

37. Svare, J. A., Schmidt, H., Hansen, B. B. & Lose, G. Bacterial vaginosis in a cohort of Danish pregnant women: prevalence and relationship with preterm delivery, low birthweight and perinatal infections. BJOG 113, 1419–1425 (2006).

38. Mercer, B. M. et al. The Preterm Prediction Study: prediction of preterm premature rupture of membranes through clinical findings and ancillary testing. The National Institute of Child Health and Human Development Maternal-Fetal Medicine Units Network. Am. J. Obstet. Gynecol. 183, 738–745 (2000).

39. Bradshaw, C. S. et al. High recurrence rates of bacterial vaginosis over the course of 12 months after oral metronidazole therapy and factors associated with recurrence. Journal of Infectious Diseases 193, 1478–1486 (2006).

40. Sivro, A. et al. Sex work is associated with increased vaginal microbiome diversity in young women from Mombasa, Kenya. J. Acquir. Immune Defic. Syndr. Publish Ahead of Print, (2020).

41. Eslami, S. et al. Lactobacillus crispatus strain SJ-3C-US induces human dendritic cells (DCs) maturation and confers an anti-inflammatory phenotype to DCs. APMIS 124, 697–710 (2016).

42. Neumann, C. et al. c-Maf-dependent Treg cell control of intestinal TH17 cells and IgA establishes host-microbiota homeostasis. Nat. Immunol. 20, 471–481 (2019).

43. Boothby, I. C., Cohen, J. N. & Rosenblum, M. D. Regulatory T cells in skin injury: At the crossroads of tolerance and tissue repair. Sci Immunol 5, eaaz9631 (2020).

44. Liebenberg, L. J. P. et al. HPV infection and the genital cytokine milieu in women at high risk of HIV acquisition. Nat Commun 10, 5227–12 (2019).

45. Masson, L. et al. Defining genital tract cytokine signatures of sexually transmitted infections and bacterial vaginosis in women at high risk of HIV infection: a cross-sectional study. Sex Transm Infect 90, 580–587 (2014).

46. Zevin, A. S. et al. Microbiome Composition and Function Drives Wound-Healing Impairment in the Female Genital Tract. PLoS Pathog. 12, e1005889 (2016).

